# Reproducibility2020: Progress and Priorities

**DOI:** 10.1101/109017

**Authors:** Leonard P. Freedman, Gautham Venugopalan, Rosann Wisman

## Abstract

The preclinical research process is a cycle of idea generation, experimentation, and reporting of results. The biomedical research community relies on the reproducibility of published discoveries to create new lines of research and to translate research findings into therapeutic applications. Since 2012, when scientists from Amgen reported that they were able to reproduce only 6 of 53 “landmark” preclinical studies, the biomedical research community began discussing the scale of the reproducibility problem and developing initiatives to address critical challenges. GBSI released the “Case for Standards” in 2013, one of the first comprehensive reports to address the rising concern of irreproducible biomedical research. Further attention was drawn to issues that limit scientific self-correction including reporting and publication bias, underpowered studies, lack of open access to methods and data, and lack of clearly defined standards and guidelines in areas such as reagent validation. To evaluate the progress made towards reproducibility since 2013, GBSI identified and examined initiatives designed to advance quality and reproducibility. Through this process, we identified key roles for funders, journals, researchers and other stakeholders and recommended actions for future progress. This paper describes our findings and conclusions.

## INTRODUCTION

### Introduction and Purpose of the Report

Preclinical biomedical research is the foundation of health care innovation. The preclinical research process is a cycle of idea generation, experimentation, and reporting of results (Figure 1).[1] The biomedical research community relies on the reproducibility of published discoveries to create new lines of research and to translate research findings into therapeutic applications. Irreproducibility limits the translatability of basic and applied research to new scientific discoveries and applications.

**Figure 1:**
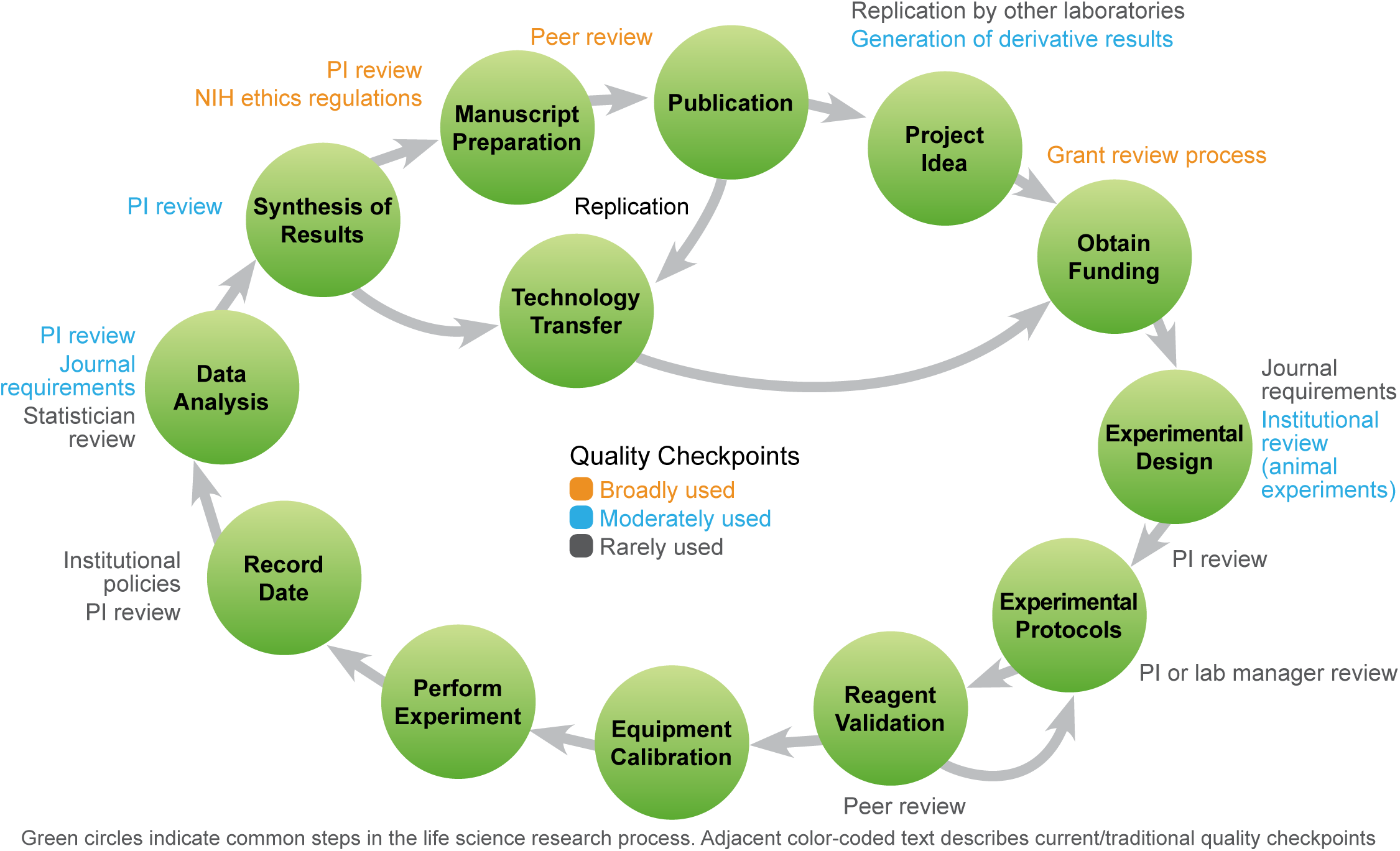
Many opportunities exist to improve reproducibility across the research life cycle. Figure from [1].

Although quality control during the research process centers on review of proposals and of completed experiments (Figure 1), opportunities to improve reproducibility exist across the entire life-cycle of the research enterprise. In fact, as Figure 1 describes, there are very few steps in the cycle where quality check points are broadly used. Recognizing these opportunities, stakeholders, such as leading scientists, journals, funders, and industry leaders are taking meaningful steps to address reproducibility throughout the research life-cycle, including commitments to scientific quality, a willingness to examine long-held research policies, and the development of new policies and procedures to improve the process of science.

The magnitude and effects of reproducibility problems are well documented. In 2012, scientists at Amgen reported that they were able to reproduce only 6 of 53 “landmark” preclinical studies.[2] GBSI released the “Case for Standards” in 2013, one of the first comprehensive reports to address the rising concern of irreproducible biomedical research. Further attention was drawn to issues that limit scientific self-correction including reporting and publication bias, underpowered studies, lack of open access to methods and data, and editorial and reviewer bias against publishing reproducibility studies (see Section IV). [3] Based on these findings, GBSI completed an economic study in 2015 and estimated that the prevalence of irreproducible preclinical research exceeds 50% with associated annual costs of approximately $28B in the United States alone.[4]

Research community stakeholders have responded to these concerns with innovation and policy. In early 2016, GBSI launched the Reproducibility2020 Initiative to leverage the momentum generated by these stakeholder-led initiatives. Reproducibility2020 is a challenge to all stakeholders in the biomedical research community to improve the quality of preclinical biological research by the year 2020. The Reproducibility2020: Progress and Priorities Report (Report), is the first to highlight progress and track important publications and actions since the issue started to get broad research community and public attention in 2013.[5, 6] The Report addresses progress in the four major components of the research process: study design and data analysis, reagents and reference materials, laboratory protocols, and reporting and review. Moreover, the Report identifies the following broad strategies as integral to the continued improvement of reproducibility in biomedical research: 1) drive quality and ensure greater accountability through strengthened journal and funder policies. 2) engage the research community in establishing community accepted standards and guidelines in specific scientific areas. 3) create high quality online training and proficiency testing and make them widely accessible. 4) enhance open access to data and methodologies.

#### Note to Reader

Terms such as reproducibility, replicability, and robustness lack consistent definition. The Report draws upon the definitions promulgated by the framework proposed by Goodman et al.[7]: “methods reproducibility” refers to the complete and transparent reporting of information required for another researcher to repeat protocols and analytical methods; “results reproducibility” refers to independent attempts to produce the same result with the same protocols (often called “replication”); and “inferential reproducibility” refers to the ability to draw the same conclusions from experimental data. The Report defines “reproducibility” to include issues affecting any of these three areas.

## Irreproducibility: Drivers and Impact

This Report is organized around key areas in the life-sciences research process where action can significantly drive improved reproducibility[4] (Figure 2):

I. Study design and data analysis
II. Reagents and reference materials
III. Laboratory protocols
IV. Reporting and review

The following sections contain detailed descriptions of each of these areas, including a review of the associated reproducibility problems, solutions, and examples of recent or current activities to promote greater quality and rigor (summarized in Table 1). The Report outlines the potential impact that lack of reproducibility has on the research community and its stakeholders (Table 2).

**Figure 2:**
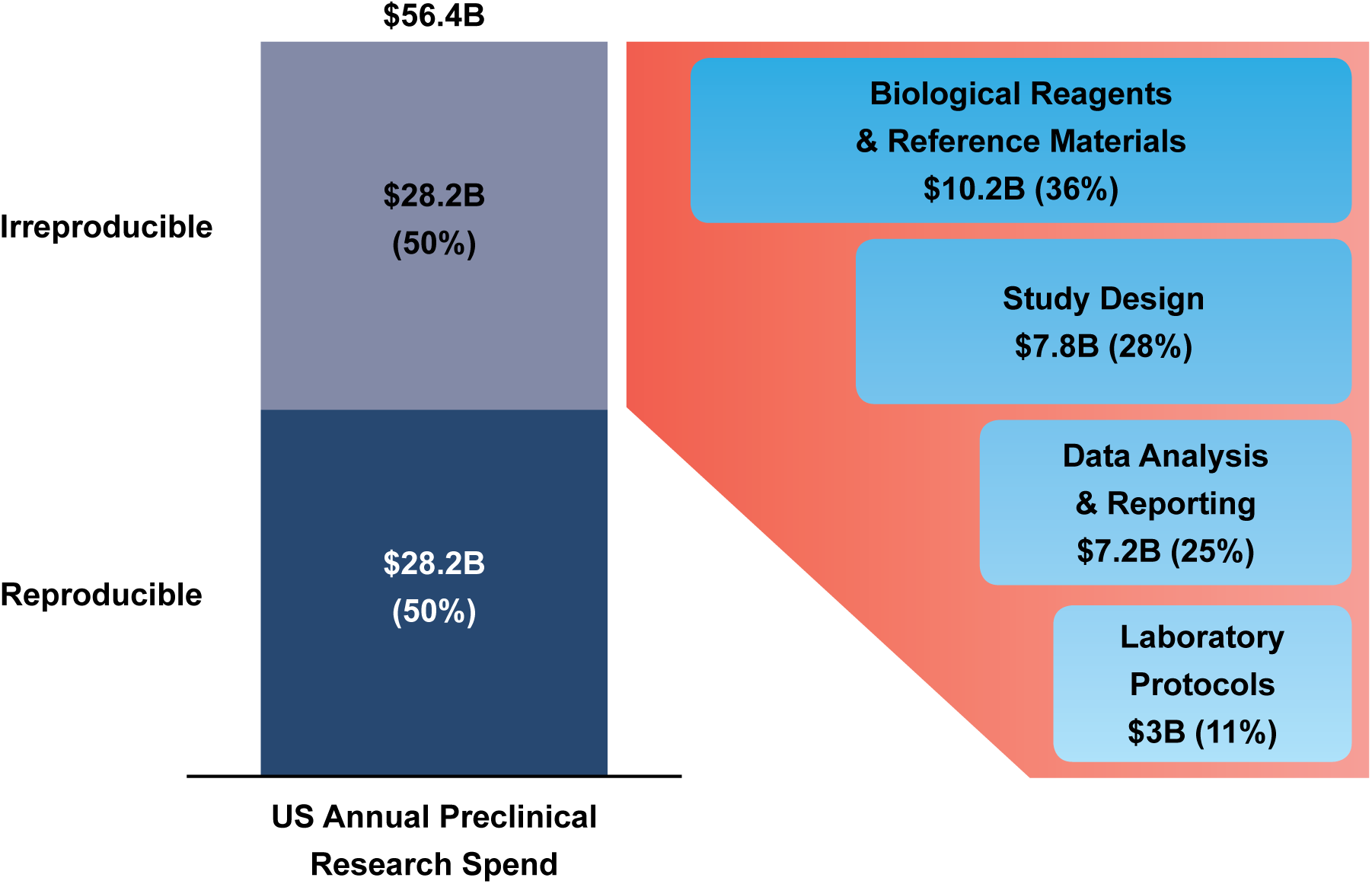
The magnitude of the reproducibility crisis and key sources of irreproducibility. Figure adapted from [4].

**Table 1:**
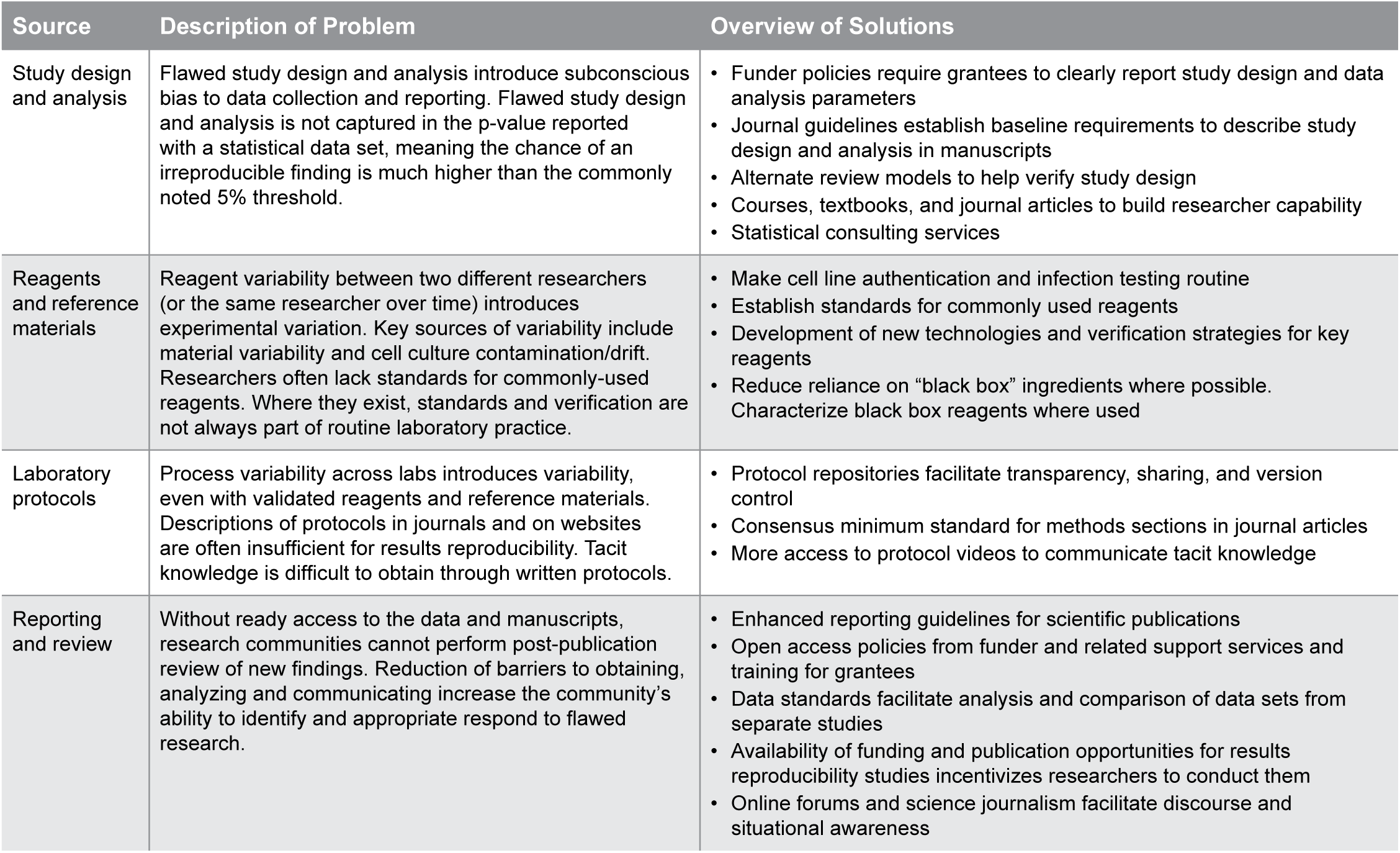
Key sources of irreproducibility and solutions

**Table 2:** Reproducibility affects all stakeholders in preclinical life sciences research

| Stakeholder | Implications of irreproducibility |
| --- | --- |
| Funders | • Impeded progress towards achieving organizational mission and goals<br>• Wasted resources spent on funding follow-on research based on a flawed premise<br>• Inefficient use of resources spent on checking, correcting, and refuting irreproducible work |
| Researchers and Research Institutions | • Adverse effect on reputation and career prospects<br>• Difficulty in obtaining future funding<br>• Failure of research projects that are based on irreproducible findings from the literature |
| Journals | • Impact of irreproducibility could negatively affect reputation, readership and journal prestige<br>• Increased administrative costs of managing retractions and errata |
| Industry | • Expensive failed clinical trials<br>• Resources wasted on failed in-house results reproduction<br>• Decreased trust in providers' products leading to decreased sales |
| Nonprofits/Scientific Societies | • Unrealized opportunities to provide value to stakeholders and members in line with organizational mission |
| Public | • Delayed realization or lost opportunities of health care benefits based on preclinical research findings, negatively impacting the discovery of life-saving therapies and cures<br>• Inefficient spending of taxpayers' money |

## METHODS

To identify key initiatives in reproducibility of biomedical research from 2013 to 2017, we conducted a review of literature, U.S. government policies, and online sources using the following keywords: reproducibility, rigor, transparency, and open access. Through these initial searches, we identified conferences on and funders of various efforts associated with reproducibility, which we used to identify other initiatives that were not identified using the keyword approach. We analyzed the information and developed recommended actions for us to promote and roles for life science stakeholders.

## RESULTS AND DISCUSSION

### I. Study design and analysis

Study design is the development of a research framework and analytical methods prior to beginning experiments. [8] A well-designed study has a research question with a rationale, and clearly defined experimental conditions, sample sizes, and analytic methods. In addition, researchers may include practices such as blinded analysis to mitigate subconscious bias. Pre-determining the research questions and sample sizes helps avoid problems such as “p-hacking” and selective reporting, where sample sizes and analytic variables are chosen based on their statistical significance rather than through a research framework (e.g., a hypothesis or an exploratory research model). However, poor study design and incorrect data analysis can sabotage even a perfectly executed experiment.

Researcher surveys suggest that study design flaws are a key source of irreproducibility. Four of the top ten irreproducibility factors identified in a researcher survey relate to poor study design and analytical procedures. [13] These findings can promote a multifaceted approach to improving study design and data analysis. Although researchers ultimately are responsible for ensuring sound study design and analysis, funder policies should encourage rigorous study design before research begins, journal requirements should facilitate better review of completed research, and training and support resources should improve researchers’ study design and analysis skills.

#### NIH study design policy

Funder policies that require good study design are especially powerful because they encourage researchers to develop rigorous study plans before beginning experimentation. Clinical research has regulatory mechanisms to review study design; for example, Phase 2 and 3 Investigational New Drug (IND) clinical trial applicants must acquire FDA approval of the study design and statistical analysis plan that includes explicit description of contingencies such as sample exclusion criteria. [14] Preclinical biomedical research is not covered by these regulatory standards, and generally has not required explicit justifications of key parameters, such as sample sizes and statistical tests, in the hypothesis and specific aims sections of proposals or in publications. For example, an analysis of 48 neuroscience meta-analyses found that 28 (57%) of the studies had a median study power of 30% or less, despite the relative ease of increasing sample size.[15] The new NIH policy (see Box 1) requires grant reviewers to explicitly incorporate several key rigor and transparency features into their peer reviews, but the policy does not add dedicated scoring line items for these areas. With respect to study design and analysis, the policy requires grant applicants to evaluate the rigor of prior studies that form the basis of a research proposal, and to justify their proposed study design. In the first round of reviews with the new guidelines, the NIH Center for Scientific Review noted that panels increasingly discussed the areas of emphasis, but that additional communication is required to get all reviewers and applicants on the same page.[16] Formal evaluations of this ongoing effort will provide valuable lessons for NIH and other funders interested in implementing their own rigor and transparency guidelines.

###### Box 1: Strengthened funder policies

As the largest and most influential research funder in the world, NIH took a major step in establishing new guidelines and going on record that NIH will address other areas where they can impact reproducibility.[9] NIH serves as an important model for other government and private research funders looking to establish greater accountability around quality and rigor.

###### NIH Rigor and Transparency Guidelines

NIH’s Rigor and Transparency Guidelines went into effect on January 25, 2016.[10, 11] This policy includes applicant and reviewer guidance in four key areas: scientific premise, scientific rigor, consideration of sex and other biological variables, and authentication of key biological and/or chemical resources. [12] Applicants are required to describe the strengths and weaknesses of prior studies cited in their scientific premise, specifically describe and justify the proposed study design, and develop authentication plans based on established standards. Because reviewers are now instructed to review applications based on these criteria, grant applicants that fail to meet the new criteria are less likely to be funded. NIH also requires grantees to report on rigor and transparency measures in their publications and the Research Performance Progress Reports submitted during the life of an award. These new guidelines underscore the need for development and propagation of study design training, pre-registration resources, and low cost authentication tools. For further information, see the NIH webpage: https://grants.nih.gov/reproducibility/index.htm

###### Box 2: Online training and proficiency testing

New approaches to training researchers should be a priority for all steps in the research cycle, including the study design training resources described in the Report. Enhanced training should be available for all levels of researchers—graduate students, post-docs, and experienced PIs. Active learning opportunities are particularly important, considering the informal apprenticeship culture of science, in which trainees learn how to design, perform, and report on their research by working with more senior scientists. However, not all senior researchers have the most current expertise or may not be able to spend the requisite time with their trainees. Surveys of researchers support this need: the 2016 Proficiency Index Assessment indicated that even experienced researchers stand to benefit from study design training, and a Figshare and Digital Science survey reported that over half of researchers wanted training on open access policies and procedures.[24, 25]

Innovative pedagogical approaches are required to ensure that training is effective and engaging for researchers at all stages of their careers. These approaches, including interactive teaching, in-lab practice, and proficiency assessments, are increasingly being explored by many institutions (see “Training and Support” example in Section I). Online training modules are a cost-effective way to provide high-quality, accessible, interactive training for researchers at all levels.

To augment these efforts, NIH has worked with the journal community to develop publication guidelines (see “Reporting and Review” below), and funded the development of researcher training programs in study design (see “Training and Support” below) as part of its Rigor and Reproducibility efforts.

#### Journal efforts to improve study design

Several studies indicate that fewer than 20% of highly-cited publications contain adequate descriptions of study design and analytic methods.[17] At least thirty-one journals have signed on to the Principles and Guidelines for Reporting Preclinical Research, which included a call for journals to include statistical analysis reporting requirements and to verify the statistical accuracy of submitted manuscripts (see “Reporting and Review” below).[18] Because these principles do not specify what these requirements should be, implementation varies by journal. One example from the *Biophysical Journal* recommends that authors consult with a statistician and requires reporting of specific information about sample sizes and statistical analyses.[19]

In the United Kingdom, the Animal Research: Reporting of *In Vivo* Experiments (ARRIVE) guidelines developed by the National Centre for the Replacement Refinement & Reduction of Animals in Research, include a checklist for researchers who perform animal studies to help researchers appropriately report study design and sample size justifications.[20] These guidelines also can be used to help ensure that researchers are planning their animal experiments correctly. As of January 2017, these reporting guidelines have been endorsed by nearly 1,000 journals and required by the major funders in the UK, including the Wellcome Trust and Medical Research Council.[21]

Some journals are prototyping alternate review models to help verify study design. As of January 2017, the Registered Reports initiative through the Center for Open Science allows selected reviewers to comment on study design and methods prior to data collection.[22] Once study design has been approved, participating journals essentially guarantee publication so long as the authors follow the study design. In addition, researchers can use the Registered Reports format to submit articles to these journals. Currently, forty-five journals are participating in this initiative. In a separate, but related initiative, the Center for Open Science’s Pre-Registration Challenge has been designed to provide training and incentives for up to 1,000 researchers to pre-register study protocols and submit manuscripts to participating journals.[23]

One journal, *Psychological Science*, currently is pilot testing statcheck software on all submitted manuscripts.[26] Statcheck and StatReviewer are tools developed by researchers to automatically review data analysis information contained in published manuscripts.[27, 28] Researchers also have deployed this tool broadly on thousands of published studies (see “Reporting and Review” below).

#### Training and support

Many life-science researchers will require training and support to satisfy the funding and publication policies described above. In the 2016 Proficiency Index Assessment (PIA), GBSI surveyed over 1,000 researchers of varying experience levels. Participants reported lower confidence in their skills in study design, data management, and analysis compared to their experimental execution skills.[24] Furthermore, research experience did not correlate with higher study design proficiency, suggesting the value of ongoing training and support in this area. New textbooks,[8, 29] online minicourses,[30, 31] and journal articles [32] can be used for course development or independent study by more senior trainees.

The positive response to study design courses established at Johns Hopkins University [33] and Harvard University [34] demonstrate the value of study design training. These courses are becoming more widespread and better tailored to the needs of life scientists, but are not universally available or required. Efforts are underway to increase the experimental design skillset of early-career students, but funding in this area has been relatively modest and in general, private funders have seen training and education as the responsibility of government funders and graduate programs. In 2014, NIH funded graduate courses on study design. Since 2014, NIH has issued a series of four funding opportunities for grantees interested in providing study design instruction for their graduate students and postdoctoral trainees through administrative supplements to existing grants.[35, 36] Several of these grantees have used the funds to develop study design training programs that are tailored to their respective research areas.[37] For more computationally-focused researchers, a Harvard course on reproducible genomics is available online for free.[38]

In addition to training, researchers now have increased access to expert support during study design and analysis. University statistics departments often provide free consulting services to affiliated researchers,[39-41] and the Center for Open Science provides a similar service.[42] The CHDI Foundation provides protocol and study design assistance, evaluation, and review to researchers studying Huntington’s disease.[43] This model may be of interest to other disease-specific funders as a low-cost investment that can improve research rigor and strengthen the community of practice in their mission area.

These training and support resources work together to improve reproducibility by increasing the general standard of rigor for all research. As researchers gain an improved understanding and awareness of study design, they can design their own studies better and more effectively communicate with statistics consultants, conduct peer review, and evaluate published findings that may inform future work.

### II. Reagents and reference materials

Reproducibility is difficult if labs are not working with the same research reagents and materials. Supplier-to-supplier variability often is poorly characterized until researchers run into problems with results reproducibility, as demonstrated by the example of synthetic albumin. The structure, stability, and immunogenicity of synthetic albumin varies across suppliers and lots, in ways that are not commonly characterized.[44] In addition, factors such as lot-to-lot material variability, cell line drift, and contamination can cause an individual researcher’s assays to change over time. Examples from other sectors suggest that these problems can be addressed with standards.

Materials developed and validated based on standards are well-characterized and demonstrate consistency. Standardized materials exhibit a predictable behavior, can be used reliably in methods reproducibility, and facilitate development of reference materials for assay validation. Standards of most well-known and often-used biological materials typically apply to particular clinical applications, such as virus strains used in influenza vaccine development.[1] Although preclinical researchers often use standardized chemical reagents (e.g., salts and sugars), few standardized biological materials exist. However, surveys suggest that life science researchers increasingly understand the need for standardized materials,[1] and the research community recently has made progress on cell line authentication and antibody validation.

#### Standards development for biomedical research reagents

Stakeholders of preclinical research include researchers, reagent manufacturers, funders, journals, standards experts, and nonprofit organizations from countries throughout the world. Recent efforts to establish antibody databases, information-sharing requirements, and international frameworks for antibody validation standards are good examples of the broad, multi-stakeholder approach required to develop consensus standards around a specific reagent

#### Good cell culture practice

One well-known example of developing standards for laboratory reagents is cell culture validation, which includes assay validation, cell line authentication, and testing for contamination.[52] Many commonly-used cell lines are available from repositories such as ATCC, as well as other nonprofit, governmental, and for-profit organizations. These organizations regularly test and validate the cells, confirming desired cell function and testing for accidental cross-contamination or infection. Researchers in two different labs can purchase validated cells from these providers and be assured that they are receiving the same product, but cells diverge once they are used in the lab. Use of shared sterile culture hoods, incubators, and reagent storage spaces can cause infection with bacteria, viruses, mold, or yeast, and result in unintentional cross-contamination of purchased cells with other cell cultures used in the lab. Even without contamination, genetic changes occur in cells through repeated culturing and experimentation, a process known as cell line drift. Despite these known problems, periodic cell line authentication and infection testing are not universally-practiced in preclinical research even though a human cell authentication standard exists.[53, 54]

As with study design, cell culture validation can be enhanced with policies from funders and journals. For example, the Prostate Cancer Foundation has been a leader in validation of cell lines used to study the disease, requiring periodic cell line authentication since 2013. NIH now requires grant applicants to describe their authentication plan as part of the Rigor and Transparency guidelines,[10] and many journals now ask researchers to perform cell line authentication.[55]

Many of the validation assays required for cell culture validation can be borrowed directly from other applications. In 2011 and 2012, ATCC organized an international group of scientists from academia, regulatory agencies, major cell repositories, government agencies, and industry to develop a standard that describes optimal cell line authentication practices, ANSI/ATCC ASN-0002-2011. The authentication assay uses Short Tandem Repeat (STR) profiling technology and is an affordable cell line authentication tool. The International Cell Line Authentication Committee’s *Database of Cross-contaminated or Misidentified Cell Lines* provides researchers with a dataset to check during the authentication process.[56] For products of animal origin, U.S. Department of Agriculture (USDA) regulations specify testing protocols for mycoplasma and select viruses [57] and test kits are commercially available.

Improving the reproducibility and translation of biomedical research using cultured cell lines must build on ongoing, multi-stakeholder efforts to raise awareness of the issues of misidentification and the role of authentication.[58] GBSI’s #authenticate campaign encourages this kind of stakeholder engagement.[59]

#### Technology and assay development

The development and propagation of standards is an iterative process. For example, recent publications highlight the simultaneous progress in cell line authentication technologies and standards development, including the establishment of reference data standards and cell line authentication policies for the broader research community.[52, 53] As technology development progresses, the standards need to be revisited and improved to reflect the current capabilities afforded by new tools.[60] For example, more affordable next generation sequencing is an increasingly useful tool to validate genome editing and characterize changes in cell behavior,[61] and mass spectrometry and lab-on-a-chip assays can help characterize sera and other liquid reagents.[62, 63]

#### Sera validation: an opportunity for standards and technology development

One opportunity to further improve cell culture validation would be to develop standards for sera production and validation. The media used to feed most cells in culture include sera, such as fetal bovine serum, that provides a variety of growth factors and other small molecules. Even authenticated cells may perform very differently in two different sera preparations. Serum is a “black box” ingredient with high variability between manufacturers and lots. Recently developed best practices include characterizing and reporting information on the particular lot(s) of serum/sera used in an experiment, and repeating an experiment with multiple lots of sera to ensure that observed phenotypes are not serum-related artifacts.[64] Serum manufacturers have begun to characterize and validate sera,[65] but no industry standard exists for reporting serum characteristics and reliability.

Further technological development could reduce reliance on sera. In serum-free culture, researchers precisely define all components of the cell culture medium rather than using a “black box” serum. Building a system with defined minimum essential components improves reproducibility and enhances scientific understanding of the key signaling molecules involved in biological processes of interest.[64] Researchers are developing and validating robust, serum-free culture systems. Clear material and validation standards are building blocks that facilitate this development.

### III. Laboratory protocols

Reproducibility requires thorough, detailed laboratory protocols. Without ready access to the original protocols, researchers may introduce process variability when attempting to reproduce the protocol in their own laboratories. The respondents of our Proficiency Index Assessment survey were more confident in their experimental skills than their study design skills. [24] Despite this relative confidence in their laboratory execution skills, researchers frequently are unable to recreate an experiment based on the experimental methods published in journals, which usually do not contain step-by-step laboratory protocols that specify every relevant variable. Further, a particular study may use a modified version of an established protocol, but state the method was “as previously described” without noting the changes. If attempts to contact authors to request the original protocols are not successful, the reader may not be able to reproduce the methods in the published work. In a *Nature* survey, nearly half of researchers felt that incomplete experimental protocol descriptions in published articles hindered methods reproduction efforts.[13] Although fewer efforts exist in this key area than in the other three areas described in this report, newly developed tools and processes designed to facilitate protocol sharing and version control may improve documentation and reduce barriers to methods reproduction.

#### Protocol repositories

Protocol repositories are an innovative approach that may facilitate transparency, protocol sharing, and version control. Researchers can upload their protocols to a repository, such as Protocols.io, precisely specifying all step-by-step instructions with links to required reagents. As the original researchers, or others, modify the protocol, they can document these changes in the repository and create their own “forked” version of the protocol. Protocols in the repository can receive a DOI number, making identification of the precise version used in a publication easier. Suppliers also can post recommended protocols for their products on these websites, which facilitates adoption of their products.

Protocol development requires a robust community of practice, so that protocols can be developed and tested by researchers in different laboratories. This practice ensures that the written instructions are understandable and replicable by a third party. Emerging on-line tools, such as BioSpecimen Commons (Arizona State University Biodesign Institute), provides a common location and uniform set of protocols and conditions for clinical sample-related SOPs. Another example is the international Protist Research to Optimize Tools in Genetics group, funded by the Gordon and Betty Moore Foundation and working on the Protocols.io website.[66, 67] As of January 2017, this group has 95 members who have contributed 31 protocols to the platform. Although this group does not focus on preclinical research, the practices established by this group are a relevant example that could be reproduced in preclinical research. Preclinical research funders may find added value with version control, protocol forking, and communities of practice in their areas of interest.

#### Improved protocol reporting in journals

The Principles and Guidelines for Reporting Preclinical Research also call for “no limit or generous limits on the length of methods sections.”[18] However, most methods sections still do not contain step-by-step protocols. Authors submitting to participating journals can include links to Protocols.io in the methods section, specifying the exact version of a protocol that was used in the study with a DOI number.[68]

Although methods journals (i.e., those dedicated to publishing detailed methods) usually provide sufficient information about protocols, most scientific publications do not. Even new techniques are not described in full detail because they build on established techniques, the methods for which are not fully described. However, some journals, such as the *Journal of Visualized Experiments*, publish original, peer-reviewed manuscripts and videos of both established and new techniques. [69] The use of videos helps to communicate technique subtleties that may not be captured in written instruction. This type of tacit knowledge often only can be obtained by visiting a laboratory and learning directly from the protocol developers.

### IV. Reporting and review

The scientific community requires ready access to publications and the original underlying data to adequately review studies and conduct results reproducibility efforts. Journal reporting guidelines improve methods reproducibility by ensuring that manuscripts contain a minimum standard of required information. Data standards further facilitate this process, as large data sets formatted in an agreed-upon, machine-readable format are easier to find, compare, and integrate across different studies. With better access to data and manuscripts, researchers now can engage in more robust post-publication review. Reducing these barriers can improve reproducibility by identifying potential flaws in published papers, making scientific self-correction and self-checking faster and cheaper.

#### Enhanced journal reporting guidelines

Journals increasingly recognize the importance of methods reproducibility and are developing more transparent and enhanced reporting guidelines. Co-led by the Nature Publishing Group, the American Association for the Advancement of Science (publisher of Science), and the NIH (as part of its Rigor and Reproducibility efforts), the scientific journal community established the Principles and Guidelines for Reporting Preclinical Research in June 2014.[18] Per the last update of the NIH website in 2016, thirty-one journals have signed on to these guidelines.[18] These guidelines provide a minimum consensus standard for statistical rigor, reporting transparency, data and material availability, and other relevant best practices, but do not specify in detail exactly what these reporting requirements should be.

More specific guidelines from journals have built upon this initial effort. These guidelines affect study design and analysis, and also are relevant to the other key areas described below. Differences in implementation of reporting guidelines may cause some short-term confusion among authors and reviewers. However, over time, implementation of these guidelines could provide long-term benefit in identifying successful approaches and best practices. Some examples of publishing guidelines include:

- Expanded reproducibility guidelines from the *Biophysical Journal* are an example of what enhanced journal guidelines look like in practice. These guidelines specifically establish reporting standards in four key areas: Rigorous Statistical Analysis, Transparency and Reproducibility, Data and Image Processing, and Materials And Data Availability.[19]
- Authors submitting to the Nature Publishing Group family of journals must complete a reporting checklist to ensure compliance with established guidelines including a requirement that authors detail if and where they are sharing their data.[70]
- STAR Methods guidelines (Structured, Transparent, and Accessible Reporting) are designed to improve reporting across Cell Press journals. These guidelines remove length restrictions on methods, provide standardized sections and reporting standards for methods sections, and ensure that authors include adequate resource and contact information.[71]
- Part of the Open Science Framework, the Transparency and Openness Promotion (TOP) guidelines for journals include template guidelines for journals interested in implementing their own reproducibility guidelines. [72] The template guidelines exist in a tiered framework, so journals can gradually implement more stringent standards as they improve their own implementation and review capability.
- Since January 2016, researchers funded by the Howard Hughes Medical Institute have been required to adhere to a set of publication guidelines that cover similar areas as the minimum consensus guidelines described above.[73]
- The Research Resource Identification Initiative establishes unique identifiers for reagents, tools, and materials used in experiments, reducing ambiguity in methods descriptions.[74]

Journals and funders can use two methods to measure and continuously improve implementation of these Guidelines: 1) stakeholder feedback studies; and 2) research measuring the frequency of compliance over time. The journal community periodically should reconvene and use data from these evaluations to identify and propagate successful implementation of the Guidelines and to update and improve the Guidelines.

#### Open access policies

Funder policies increasingly mandate access to data and publications (Box 3). As of October 2016, sixteen U.S. government funding agencies require their grantees’ publications to be open access within a year of the publication date, and thirteen of these funders, including the NIH, require data management plans to be included in research proposals.[75] Globally, the online research repository Figshare predicts that by 2020, all funders in the developed world will require openness.[76]

###### Box 3: Improved reagent standards: the Antibody Initiative

The research community has acknowledged that antibodies are an area of widespread error and inaccuracy.[45] The Antibody Validation Initiative, involving stakeholders throughout the research community and led by GBSI, is an example that could be replicated in other scientific areas (e.g. both stem cells and synthetic biology are areas where a greater emphasis on development of standards and best practices are needed to ensure quality and advance discovery). Antibodies are key reagents in preclinical research for activities as diverse as protein visualization, protein quantification, and biochemical signal disruption. Antibody performance is variable, with differences in specificity, reliability, and functionality for different types of experiments (e.g., Western blotting and immunofluorescence), manufacturers, and lots, harming reproducibility.[46] Stakeholder solutions include antibody databases such as the CiteAB database [47] and repositories such as the proposed universal library recombinant antibodies for all human gene products.[48] In all cases, validation is a key component of the solution.

NIH specifically highlights antibody authentication in the Rigor and Transparency guidelines,[10] providing additional impetus for new standards, policies, and practices. Researchers, manufacturers, pharmaceutical companies, funders, and journals have held dedicated conferences on antibody validation (e.g. [49]). In 2016, the International Working Group on Antibody Validation (IWGAV) qualitatively identified key validation “pillars” that may be suitable for assessing antibody performance.[50] Seeking to build on the IWGAV recommendations, GBSI and The Antibody Society organized a workshop for all stakeholder groups to develop actionable recommendations to improve antibody validation.[51] Stakeholder groups recognized the shared responsibility of antibody validation and effective communication of validation methodology and results. In addition, they highlighted the need for continued, multi-sectoral engagement during the development of standards for validation, which may vary by use case, and information-sharing, which may vary by stakeholder.

Since the workshop, GBSI established seven multi-stakeholder working groups to draft validation guidelines for the major antibody applications. Validation guidelines will include an application-specific point system to quantify antibody specificity, sensitivity, and technical performance. The Antibody Validation Initiative also includes a Producer Consortium to address issues of common concern for producers and a Training and Proficiency Assessment program to ensure the highest quality of validation.

###### Box 4: Enhanced open access to data and methodologies

Both governmental and private funders have undertaken significant policy changes to mandate open access to data sets and publications. Funders are generally moving towards more open access, mandating or encouraging researchers to publish in open access journals, paying open access fees, and requiring manuscript archival when researchers publish in more restrictive journals.

Large funders are leading the drive towards open access. NIH spends roughly $4.5 million on PubMedCentral, [85] and requires all grantees to deposit articles and/or manuscripts in this open repository within twelve months of publication.[86] The Gates Foundation and HHMI have leveraged the NIH’s investment by requiring their own grantees to archive manuscripts in PubMed.[78, 87] Gates has gone one step further on open access, requiring all publications to be immediately available in open access “Gold” format.[78] The Gates Foundation has also developed tools to assist its grantees with compliance with these new open access policies.[83]

As major funders increasingly mandate open access, more journals are providing open access options for authors. Many journals provide Creative Commons copyright options, providing a uniform set of standards. The increased adoption of Creative Commons licenses by journals, especially unrestricted CC-BY licenses, reduces the barrier to adoption of open and transparent sharing permissions.[82]

Private funders have taken a variety of approaches to promoting open access, such as increasingly requiring either full open access or archived manuscripts as a condition of continued funding (reference [77] contains a summary of many institutions’ policies). The Bill & Melinda Gates Foundation is a leader among philanthropic organizations in formulating and implementing open access policies. Beginning in January 2017, the Gates Foundation’s Open Access Policy requires immediate open access (“Gold” access) for all publications and underlying data generated by authors that it supports. [78] Many journals already have open access options that comply with the Gates Foundation policy, but some high-profile journals such as *Nature*, and *Science*, did not have Gates-compliant policies as of January 2017.[79] In response to this policy change, AAAS, the publishers of *Science*, reached a provisional agreement with the Gates Foundation to make Gates-funded publications in AAAS journals open access.[80] Similarly, the *Cell Press* family of journals has special agreements with a number of funders, including Gates, that allow immediate open access for a fee.[81] This issue warrants further attention as funders and journals continue to negotiate around access permissions. The Wellcome Trust has a similar policy, encouraging immediate open access but allowing a six month delay. Both the Wellcome Trust and Gates Foundation provide dedicated funding to support open access fees imposed by journals where appropriate, and prefer the unrestricted Creative Commons-BY license.[82]

While this represents real progress, these policies can be a source of confusion for researchers. In a recent survey of over 1,000 researchers by Figshare and Digital Science, 64% of researchers who have made their data open could not recall what licensing rights they had granted on the data (e.g. CC-BY, CC-BY-NC).[25] Additionally, 20% of researchers were unaware whether their funders had an open data policy and most researchers welcomed additional guidance on their funders’ openness policies,[25] suggesting the need for increased education and support. One facet of the Gates Foundation solution to this problem is a new service called Chronos. The Chronos service guides users through submission to services that are compliant with Gates’ policy, automatically pays open access fees, and archives manuscripts on PubMed.[83] The Gates Foundation expects to scale Chronos to additional funding organizations.[84]

The leadership of funders has led several journals to allow authors to self-archive manuscripts on preprint servers such as arXiv or bioRxiv before publication. Some journals, such as *Nature*, *F1000 Research*, and *PeerJ*, also have their own pre-print option.[88] PubMed Central and European PubMed Central also provide open full text archives. The precedent set by these large funders has established an infrastructure and leadership base that smaller funders may be able to leverage in the development and advancement of their own open access policies. Supported by the Laura and John Arnold Foundation, the Center for Open Science also has developed implementation guidelines for funders interested in establishing transparency and openness policies. [72] Like the journal guidelines, the TOP funder policies are tiered to allow funders to implement more stringent standards over time.

#### Data standards

Policies that ensure open access to the original underlying data and materials can be leveraged more effectively when the data from different studies can be compared easily. Common standards have been incorporated into reporting policies for journals. For example, the Addgene Vector Database provides a repository of published and commercially-available expression vectors.[89] At least thirty-one journals recommend or require authors to submit their plasmids to the Addgene repository.[90] Addgene performs sequencing to verify submission quality,[91] and requires each contributor to provide the same types of information in a uniform format, making the database easily searchable and comparable.

The Addgene approach works well for plasmids, which consist of a relatively limited number and size compared to high-throughput, whole genome sequencing data sets. As next generation techniques become more widespread, data standards will become even more important. These data standards include metadata (i.e., information about the data set), data fields, and file formats. With data standards, large data sets become much easier to download and interpret, because users do not have to spend valuable and expensive computational time modifying existing analysis tools to fit each new data set. Researchers have proposed a series of metadata checklists for high-throughput studies.[92] Similar to the development of reagent standards described above, updated data standards will require multi-stakeholder collaboration within the community of practice, harnessing existing standards where possible and harmonizing divergent practices where appropriate.

#### Post-publication review

Scientific review is an ongoing process that continues well after peer-review and publication. The broader scientific community may identify issues that were not highlighted by the peer reviewers, and other researchers may attempt to reproduce a study on their own. Because the post-publication review process may require experimentation, it warrants dedicated resources.

Despite the time commitment and added value to science, the research community typically does not reward post-publication review. Historically, funding agencies and tenure boards do not tend to reward results reproducibility studies, and researchers can have trouble convincing journals to review and accept such manuscripts. However, stakeholders from different sectors now are dedicating resources to results reproduction. The Laura and John Arnold Foundation currently is funding a cancer biology results reproducibility study as part of its Reproducibility Project series. The first five attempts to reproduce papers as part of this effort were published in January 2017 in the journal *eLife*, an open access journal supported by the Howard Hughes Medical Institute, Max Planck Gesellschaft, and the Wellcome Trust.[93] Two of these five studies successfully reproduced the original findings, one study did not, and two attempts were inconclusive. Because the project seeks to reproduce approximately 50 papers, conclusions about the Project’s reproducibility rates at this early stage (i.e., after five experiments) would be premature. An earlier project, Reproducibility Project: Psychology, attempted to reproduce 100 original psychology findings, successfully reproducing one-third to one-half of the results.[94] Another open access publication, *F1000Research*, established the *Preclinical Reproducibility and Robustness Channel* as a platform dedicated to reproducibility of published papers.[95]

Researchers attempting to raise concerns to editors about irreproducible or incorrectly analyzed results found in published articles describe many barriers to the process of raising these concerns, including lack of clarity and transparency from journals in the post-publication review process.[96] Similarly, journals do not always have a clearly-defined retraction process that mirrors the submission and peer review processes. Much like the stakeholder discussions on study design, cell line authentication, and open access, the retraction process is an important topic that warrants engagement by the research community. The Committee on Publication Ethics has established best practices for Retraction Guidelines,[97] which may provide an opportunity for this discussion.

Websites like PubMed Commons and PubPeer provide an informal mechanism to facilitate post-publication review and results reproduction attempts by providing a discussion forum for researchers to openly discuss scientific publications. Discussions on these platforms can occur much faster than the pace of published technical commentaries in journals and provide opportunities for more scientists to contribute. Last year, researchers undertook a widespread deployment of the automated statcheck algorithm on nearly 700,000 experiments from over 50,000 papers, and automatically generated comments on PubPeer for each paper.[98] This automated tool helps researchers identify papers that deserve further review and discussion about solutions, such as retraction or publication of counter studies. Discussions on open blogs are a double-edge sword. Whereas rapid turnaround and informal discussion can stimulate productive scientific debate, unmoderated discussion can also lead to unwarranted criticism of legitimate studies. In contrast, technical commentary in journals is refereed by an editor who can help organize and moderate the discussion.

The sheer volume of published research increases the difficulty of identifying and tracking publication errors. Science journalism is another tool that can improve reproducibility. Science reporters, such as the authors of Retraction Watch,[99] bring publicity to reproducibility and retraction news, which can galvanize the scientific community to action. For example, replicability of the initial paper describing the NgAgo genome editing technique has been the subject of fierce debate in the community wherein researchers described their difficulties in reproducing the paper’s claims on internet and scientific news sites. The technique drew so much attention that over 100 researchers attempted to reproduce the technique in the first few months after publication, but less than 10% were successful.[100] The controversy resulted in three peer-reviewed publications, all of which documented a failure to reproduce the original study, and researchers now are trying to understand the reasons for irreproducibility.[101]

Retraction Watch also partners with the Center for Open Science to generate a database of retractions, as some retracted articles still are cited frequently after retraction.[102] Researchers armed with this database can avoid using retracted work as a (shaky) foundation for new studies, thereby increasing their chance of success. By reading about reproducibility and retraction news, researchers can learn about the common pitfalls that can cause retractions and new resources available to help them improve the reproducibility of their work, such as the initiatives described in this report. However, highly-visible retractions are a potential threat to public confidence and support for science, as the lay public reads more about retractions and irreproducibility. This further highlights the urgent need for the scientific community to act on the initiatives described in this report and make meaningful improvements to reproducibility.

## Conclusion: A Path Forward

Irreproducibility is a serious and costly problem in the life sciences. Measured reproducibility rates are shockingly low, requiring significant effort to solve this problem. Many stakeholders now recognize the importance of reproducibility and are taking steps to develop and implement meaningful policies, practices, and resources to address the underlying issues. The lessons learned from these early efforts will assist all stakeholders seeking to scale up or replicate successful initiatives. The research community is making progress to improve research quality. By prioritizing the strategies outlined in the Report, stakeholders in life science research will continue to make progress in improving reproducibility and in turn have a profound positive impact on the subsequent development of treatments and cures.

However, the authors would be remiss if we ignored a transcending challenge facing the research community and their willingness to voluntarily accept these positive steps in addressing reproducibility: the current rewards system in academia, including constant pressure to obtain grants and publish in “high impact” journals. The research culture, particularly at academic institutions, must seek greater balance between the pressures of career advancement and advancing rigorous research through standards and best practices. We believe that the many initiatives described in this Report add needed momentum to this emerging culture shift in science, but additional leadership and community-wide support will be needed to better align incentives with reproducible science and effect this change.

Continued transparent, international, multi-stakeholder engagement is the way forward to better, more impactful science. GBSI calls on all stakeholders – individuals and organizations alike – to take action to improve reproducibility in the preclinical life sciences by joining an existing effort, replicating successful policies and practices, providing resources to results reproduction efforts, and/or taking on new opportunities. Table 3 contains specific actions that each stakeholder group can take to enhance reproducibility.

In its leadership role, GBSI will:

- work with journals and funders to encourage policies that increase rigor, accountability and open access to data and methodologies,
- lead the effort toward improving the validation of reagents—particularly cells and antibodies— and work with the research community to explore other scientific areas (e.g. stem cells and synthetic biology) where a greater emphasis on development of standards and best practices are needed to ensure quality and advance discovery,
- ensure high quality, accessible online training modules available to both emerging and experienced researchers who are eager to improve their proficiencies in new and evolving best practices, and
- continue to track reproducibility efforts through the Reproducibility2020 Initiative.

The preclinical research community is full of talented, motivated people who care deeply about producing high-quality science. We are optimistic about the potential to improve reproducibility, and look forward to contributing to the effort.

**Table 3:** Reproducibility2020 action plan

| Stakeholder | Actions to Improve Reproducibility in Preclinical Research |
| --- | --- |
| Funders | • Enact policies requiring study design pre-registration, cell line authentication and reagent validation, laboratory protocol transparency, and open access to publications. Provide relevant funding commitments where necessary<br>• Include specific line items in grant review to score reproducibility factors<br>• Provide resources for study design training and statistics consultation for grantees and grant applicants<br>• Fund the development of open access and transparency tools, and additional research to better characterize reproducibility<br>• Fund the development of new technologies and methods that enhance reproducibility<br>• Encourage grantees to develop communities of practice for protocol sharing and testing, and dedicate resources to facilitate and incentivize these communities<br>• Fund innovative training programs including online modules |
| Researchers and Research Institutions | • Make online accessible training modules available that address all major components and evolving approaches of the research process<br>• Explore new approaches to mentorship and accountability to ensure that emerging researchers (i.e., graduate students and postdocs) receive necessary training and supervision from experienced PIs<br>• Implement lab policies that improve reproducibility, such as reagent validation and documentation, routine cell line authentication, and independent reproduction of results by another researcher in the lab<br>• Develop institutional policies and an organizational culture that values and rewards reproduction studies, study design pre-registration, protocol sharing, and open access<br>• Organize online communities of practice to facilitate discussion and sharing of information within the field<br>• Participate in multi-stakeholder groups that develop reproducibility policies and guidelines<br>• Explicitly consider reproducibility issues during peer review of grants and manuscripts<br>• Develop new technologies and methods that improve reproducibility and assist in validation and authentication processes<br>• Explore new technologies including lab/bench automation and robotics to ensure greater precision and minimize errors<br>• Perform results reproduction studies and publish the results<br>• Explore new incentive structures for career advancement that move away from the traditional impact factor and funding paradigms to reward greater data and methods transparency, adherence to best practices and standards, and reproducibility of published work |
| Journals | • Adopt more stringent reporting and transparency guidelines, such as TOP Level 3<br>• Provide cost-effective open access publication options under CC-BY licenses<br>• Require cell line authentication and promote antibody validation guidelines, as they become available.<br>• Allow archiving of submitted manuscripts before publication<br>• Publish reproduction studies and technical commentary<br>• Consider pre-registered review models that enable rigorous peer review of study design<br>• Encourage greater use of pre-print platforms<br>• Work with researchers to establish data and metadata standards for reporting (e.g., next-generation sequencing)<br>• Require authors to link to version-controlled protocols<br>• Conduct surveys of researchers to better understand reproducibility issues and obtain feedback on journal guidelines and policies<br>• Report on reproducibility issues in the editorial and news section of the journal |
| Industry | • Transparently communicate the results of in-house replication attempts<br>• Enhance protocol transparency, discussion, and version control, especially for reagents and kits<br>• Provide validation data and technical support for reagents and kits<br>• Participate in the establishment of materials standards |
| Nonprofits/Scientific Societies | • Convene multidisciplinary groups to establish relevant standards, including materials standards for commonly-used reagents, and data standards for commonly-used experimental methods<br>• Provide professional development for researchers to improve research proficiencies, particularly in the areas of study design, data analysis, reagent validation, and reporting transparency<br>• Convene meetings focused on reproducibility to facilitate sharing of best practices and develop new policies and procedures |
| Public | • Stay aware of reproducibility news to promote a culture of accountability |

## Acknowledgments

The authors would like to acknowledge Dr. Kavita Berger and Ms. Allison Mistry from Gryphon Scientific for their review of the manuscript.

